# Structure and function of the peatland prokaryotic microbiome

**DOI:** 10.1101/2025.08.21.671631

**Authors:** Simon Man Kit Cheung, Richard D. Pancost, Mike Vreeken, Angela V. Gallego-Sala, Casey Bryce

**Affiliations:** School of Earth Sciences, Cabot Institute for the Environment, University of Bristol, Bristol, UK; Organic Geochemistry Unit, School of Chemistry, University of Bristol, Bristol, UK; Department of Geography, Faculty of Environment, Science and Economy, University of Exeter, Exeter, UK

**Keywords:** Microbial diversity, metagenome, wetland, tropical peatland, incomplete denitrification, climate change

## Abstract

Peatlands are vital long-term carbon sinks but also sources of trace greenhouse gases, with the balance largely governed by microbial processes. However, the structure and function of peatland microbial communities are currently poorly characterised beyond the local scale, limiting our understanding of their response to climate change. Here, we present a large-scale analysis of the peatland prokaryotic microbiome by leveraging a curated dataset of 109 publicly available metagenomes from 20 near-natural peatland sites worldwide. Despite limitations in the publicly available data, including geographic coverage and sampling depth, our analysis indicates that dominant vegetation and peatland type are the strongest predictors of peatland microbiome taxonomy and function. We show that obligate hydrogenotrophic methanogens of the family Bog-38, previously reported to be prevalent in diverse high methane-flux habitats, are abundant in northern peatlands but occur only at low abundance in tropical peatlands, suggesting a limited contribution to methane cycling in these ecosystems. In contrast to northern peatlands, the tropical peatland microbiome exhibits a broad metabolic potential for multiple key peatland biogeochemical processes, including methanogenesis and denitrification, across all depths. Intriguingly, we observe an impaired potential for complete denitrification in a significant proportion of the northern fen metagenomes, with potential implications for nitrous oxide emissions as a greenhouse gas. Overall, our study uncovers large-scale patterns and ecological drivers of the peatland prokaryotic microbiome based on existing metagenomic data, enhancing our understanding of its response to climate change and highlighting major data gaps for future research.

## Introduction

Peatlands cover only ∼3% of the Earth’s land area but are estimated to store up to one third of terrestrial organic carbon (C)[1]. Northern mid- and high-latitude peatlands account for over 90% of global peatland area and have sequestered over 500 Gt C[2]. These peatlands are widely classified as ombrotrophic (rainwater-fed) bogs and minerotrophic (groundwater-fed) fens[3], although several other peatland classification systems also exist[4]. Tropical peatlands do not fall neatly into that classification, although lowland domed bogs as well as highland bogs and fens also exist in the tropics. Tropical peatlands have a much lower geographic coverage but are estimated to store 152–288 Gt C – up to approximately one-third of the C stored by global peatlands[5]. Peatlands are important long-term carbon sinks but can also be natural sources of greenhouse gases such as methane (CH_4_) and nitrous oxide (N_2_O)[6]. This balance depends on microbial processes in peatlands, which are in turn affected by environmental factors and climate conditions[7]. For instance, climate warming could lower the peatland water table via increased evapotranspiration, thereby reducing anaerobic conditions and CH_4_ emissions, while enhancing aerobic respiration and carbon dioxide (CO_2_) release[6, 7].

Shotgun metagenomics, in which all the DNA in a sample is sequenced and analysed, allows profiling of the taxonomic composition and metabolic potential of microbial communities (microbiomes) in the environment[8]. Peatland metagenomic studies have been conducted in bogs and fens across Europe[9–11], North America[12–14], and Asia[15–17], and more recently, in tropical peatlands[18, 19]. These studies have provided important insights into the role of the peatland microbiome in the cycling of C[9, 11, 14, 16], nitrogen[15], and sulphur[10], as well as others, such as methylmercury production[13]. However, to fully understand ecosystem functioning, it is necessary to study large-scale distribution patterns and functional gene repertoires of the microbial inhabitants[20]. To date, the structure, function, and ecological drivers of the peatland microbiome have not been examined beyond the local scale, restricting our understanding of its response to large-scale environmental variation and climate change. In this study, we conducted an analysis of the peatland prokaryotic microbiome by leveraging a dataset of 109 publicly available metagenomes from 20 near-natural peatland sites worldwide. Our objectives were to (1) identify the large-scale patterns in the structure and function of the peatland prokaryotic microbiome, (2) elucidate the main ecological drivers of peatland microbiome taxonomy and function, and (3) identify gaps in our understanding due to incomplete data coverage.

## Results and discussion

### A large collection of metagenomes enables large-scale meta-analysis

We compiled a large dataset comprising 109 publicly available peatland metagenomes collected from 20 near-natural peatland sites across four continents: North America, South America, Europe, and Asia (Fig. 1, Supplementary Table 1). This dataset included six ombrotrophic bogs (n = 36) and nine minerotrophic fens (n = 44) from the mid- and high latitudes of the Northern Hemisphere, as well as five lowland tropical peatlands (TP) (n = 29). These peatlands span a latitudinal range of 74° (6° S to 68° N) and an elevation range from 11 m to over 3600 m. The northern bogs encompassed both lowland (n = 23) and alpine settings (n = 13) (pH: 3.8–5.9; four sites with available data); the northern fens comprised nutrient-rich and nutrient-poor systems (pH: 4.1–7.4; seven sites), although for some sites nutrient data were unavailable; and the TPs included both forested (n = 23) and herbaceous vegetation types (n = 6) (pH: 2.5–5.6; four sites). Although TPs exhibit a significant diversity[21], they were treated as a single category in this study due to a lack of data on different peatland subtypes. The median water table levels were −10.5 cm (range: −11 to 3 cm; three sites), 0 cm (−13.1 to 23 cm; five sites), and −5 cm (−10 to 0 cm; four sites) for bogs, fens, and TPs, respectively.

**Fig. 1.**
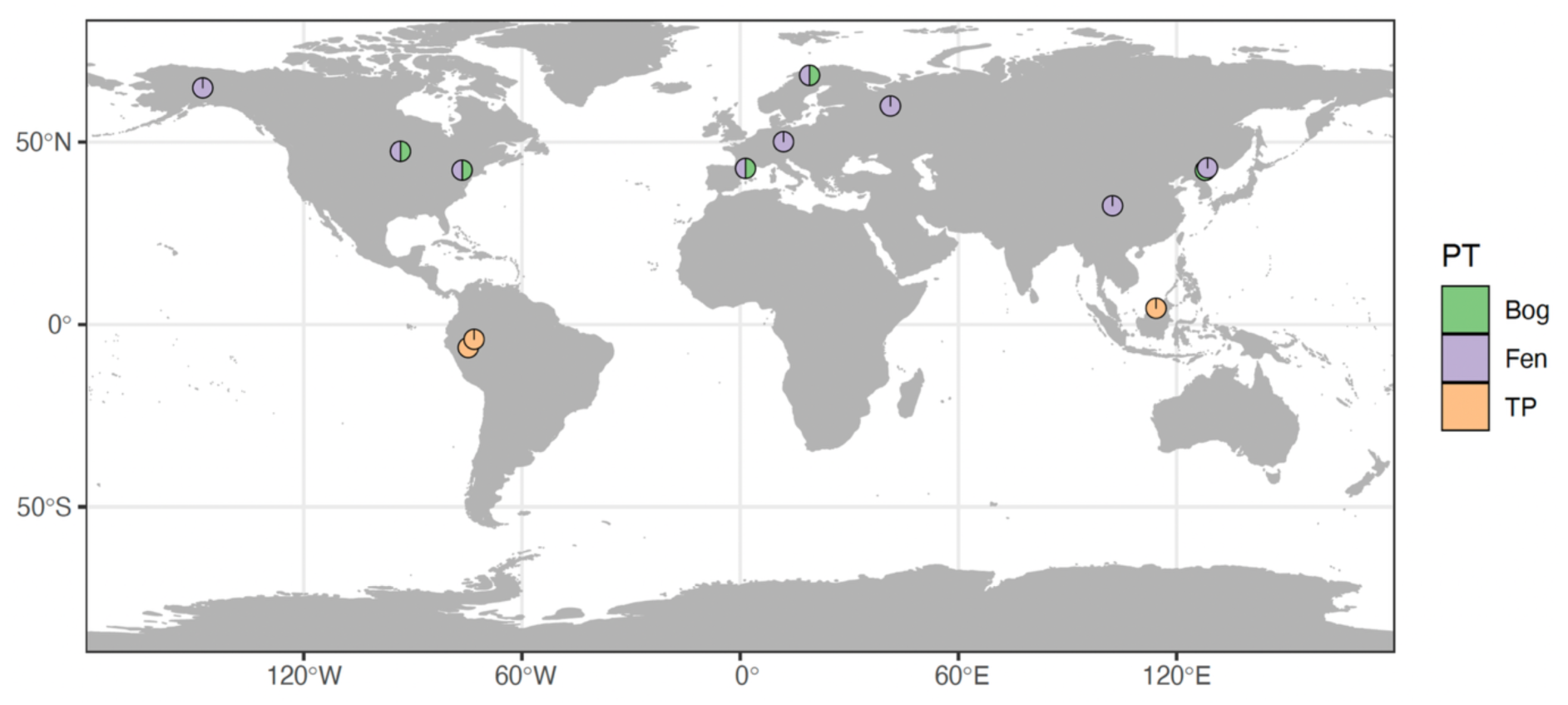
Geographic distribution of peatland metagenomes included in this study. A total of 109 metagenomes were collected from six northern bogs (n = 36), nine northern fens (n = 44), and five tropical peatlands (TP) (n = 29). The peatland types (PT) included in each location are indicated by the colour of the pies. For bog and TP, the number of pies shown on the map is smaller than the actual number of sites due to their close proximity.

The dataset included samples collected from 2.5 cm to 195 cm in depth, although most of them were from relatively shallow depths (0–10 cm: n = 48, 10–20 cm: n = 22, 20–50 cm: n = 21, below 50 cm: n = 18). It should be noted that because water table levels vary among peatland types and sites, samples collected from similar depths may experience markedly different hydrological conditions, with the water table separating oxic and anoxic peat layers. Consequently, neither depth groups nor absolute depth necessarily represent equivalent environmental conditions across sites, and direct comparisons should therefore be interpreted with caution. The distribution of samples across depth groups was uneven among peatland types. For instance, 76% of the TP samples were collected from the upper 20 cm of the peat profiles compared to 56% in bogs. At the time of this analysis, no metagenomes from peatlands located on the African or Australian continents, or below 200 cm in depth were publicly available. In addition, several of the world’s largest peatland complexes were not represented, including the Congo Basin in Central Africa[22] and the Hudson Bay Lowlands in Canada[23]. Consequently, the currently available metagenomic resources, and therefore our dataset, do not fully represent the global peatland landscape. As in any meta-analysis, variation in experimental protocols, such as DNA extraction methods and sequencing platforms, among the included studies may raise concerns about batch effects. However, such effects are generally considered minimal when the effect size of the variable of interest is large[24].

### Peatland type and peat depth shape the taxonomic and functional diversity of the peatland prokaryotic microbiome

We first examined the effects of peatland type and other environmental variables on the alpha diversity (diversity within a sample) and beta diversity (diversity between samples) of peatland microbiome taxonomy and function. Results showed that peatland type significantly influenced the taxonomic and functional alpha diversity of the peatland microbiome, as assessed by the Shannon diversity index (Kruskal– Wallis test, *p* < 0.001) (Fig. 2A, D). The northern bog microbiome exhibited a lower taxonomic and functional alpha diversity compared with the northern fen and TP microbiomes (Dunn’s test, *p* < 0.05). Taxonomic alpha diversity in northern fens was comparable to that in TPs; however, the TP microbiome was functionally more diverse than northern fens (*p* < 0.05). The lower taxonomic alpha diversity observed in northern bogs agrees with previous research comparing bog and fen sites of a mountainous peatland[11], and is likely a result of environmental filtering due to acidic and nutrient-poor conditions in bogs.

**Fig. 2.**
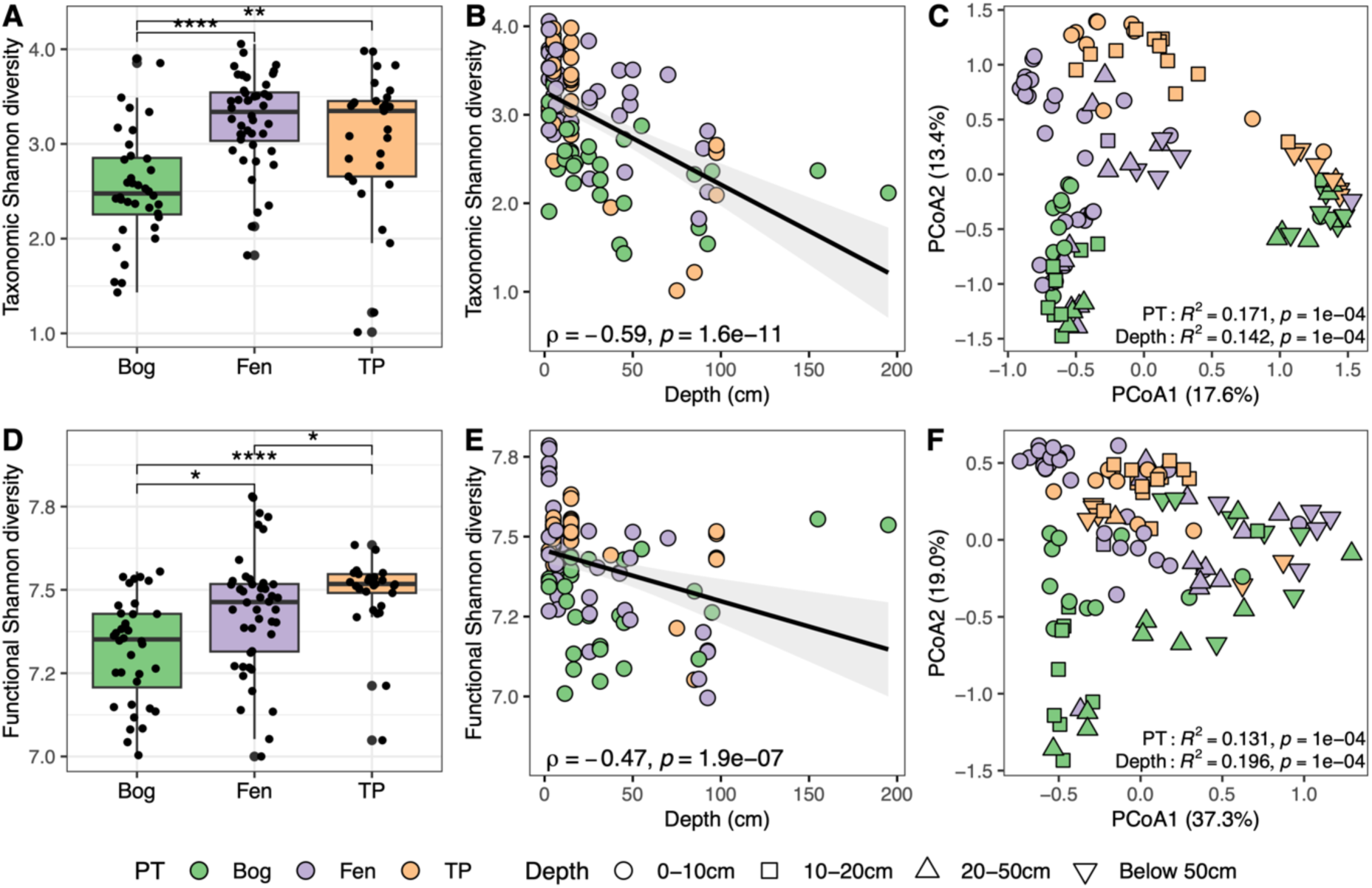
Peatland type and depth determine the taxonomic and functional diversity of the peatland microbiome. Scatter box plots of **(A)** taxonomic and **(D)** functional alpha diversity among peatland types (PT) based on the Shannon diversity index. Centre lines indicate the medians, box limits indicate the upper and lower quartiles, and whiskers extend to 1.5x interquartile range. Spearman’s correlations between depth and **(B)** taxonomic and **(E)** functional Shannon diversity. Grey areas around the regression lines represent regions of 95% confidence interval. Principal coordinates analysis (PCoA) plots of **(C)** taxonomic and **(F)** functional beta diversity based on the Bray–Curtis (abundance) dissimilarity metric. Values on the axes represent the percent variations explained. PERMANOVA statistics are provided on the plots. In each panel, each dot represents an individual sample (n = 109). All panels share the same colour scheme for PT, whereas panels C and F share the same annotation for depth. TP, tropical peatland. \**p* < 0.05, \*\**p* < 0.01, \*\*\**p* < 0.001, \*\*\*\**p* < 0.0001.

We observed moderately strong negative correlations between peat depth and both taxonomic (Spearman’s ρ = −0.59, *p* < 0.001) and functional alpha diversity (ρ = −0.47, *p* < 0.001) of the peatland microbiome across all peatland types (Fig. 2B, E). Depth is closely related with the physicochemical properties of peat and serves here as a proxy for the shifting redox and oxygenation gradients along the peat profile[3]. A reduced microbial alpha diversity along peat depth has also been reported in studies based on the 16S rRNA gene[17], and may be attributed to a more selective, low oxygen environment with poor organic matter (OM) quality at greater depth.

Peatland type and depth also influenced the taxonomic and functional beta diversity of the peatland microbiome, as assessed by both the abundance-based Bray–Curtis and presence/absence-based Jaccard distance metrics (PERMANOVA, *p* < 0.001) (Fig. 2C, F, Supplementary Fig. 1). Notably, while peatland type exhibited a larger effect size (R^2^) than depth on taxonomic beta diversity, depth, in contrast, showed a larger effect size on functional beta diversity.

Among the other continuous variables, pH and net primary productivity (NPP), an indicator of organic carbon availability, were found to be positively, albeit weakly, correlated with taxonomic and functional alpha diversity, respectively (ρ ∼ 0.33, *p* < 0.05) (Supplementary Fig. 2, 3). Interestingly, a latitudinal diversity gradient was observed in peatland function (ρ = −0.48, *p* < 0.001), but not in taxa (*p* > 0.05) (Supplementary Fig. 4).

### Dominant vegetation and peatland type are the strongest predictors of peatland microbiome taxonomy and function

We then sought to identify the ecological drivers of peatland microbiome taxonomy and function using variation and hierarchical partitioning, an approach which calculates the relative importance of individual predictors based on their combined unique and shared effects across all model subsets[25], based on a subset of metagenomes with associated pH data (n = 88). Dominant vegetation and peatland type emerged as the strongest predictors of both peatland microbiome taxonomy and function, explaining more than 8.2% and 6.0% of the variation, respectively (Fig. 3). These findings corroborate previous local studies, which show that changes in peatland vegetation composition alter microbial community composition and function[26, 27]. Vegetation cover type has also been shown to exert significant impacts on the microbial community structure in soil at the continental-scale[28]. As expected, a significant proportion of the overall importance of these umbrella variables was attributed to their average shared effects with other predictors. For function in particular, this reached ≥ 71.0% based on the Bray–Curtis dissimilarity metric, indicating that most of their effect was due to correlations among the involved predictors (Supplementary Table 2). However, their high unique effects indicate that they also accounted for substantial variation not explained by the other variables. Other potentially important physico-chemical variables, including moisture content, organic matter content, and nutrient concentrations[18], were not consistently reported across the studies included here and therefore were not examined in the current analysis. However, their effects are presumably partially accounted for by depth and peatland type, respectively.

**Fig. 3.**
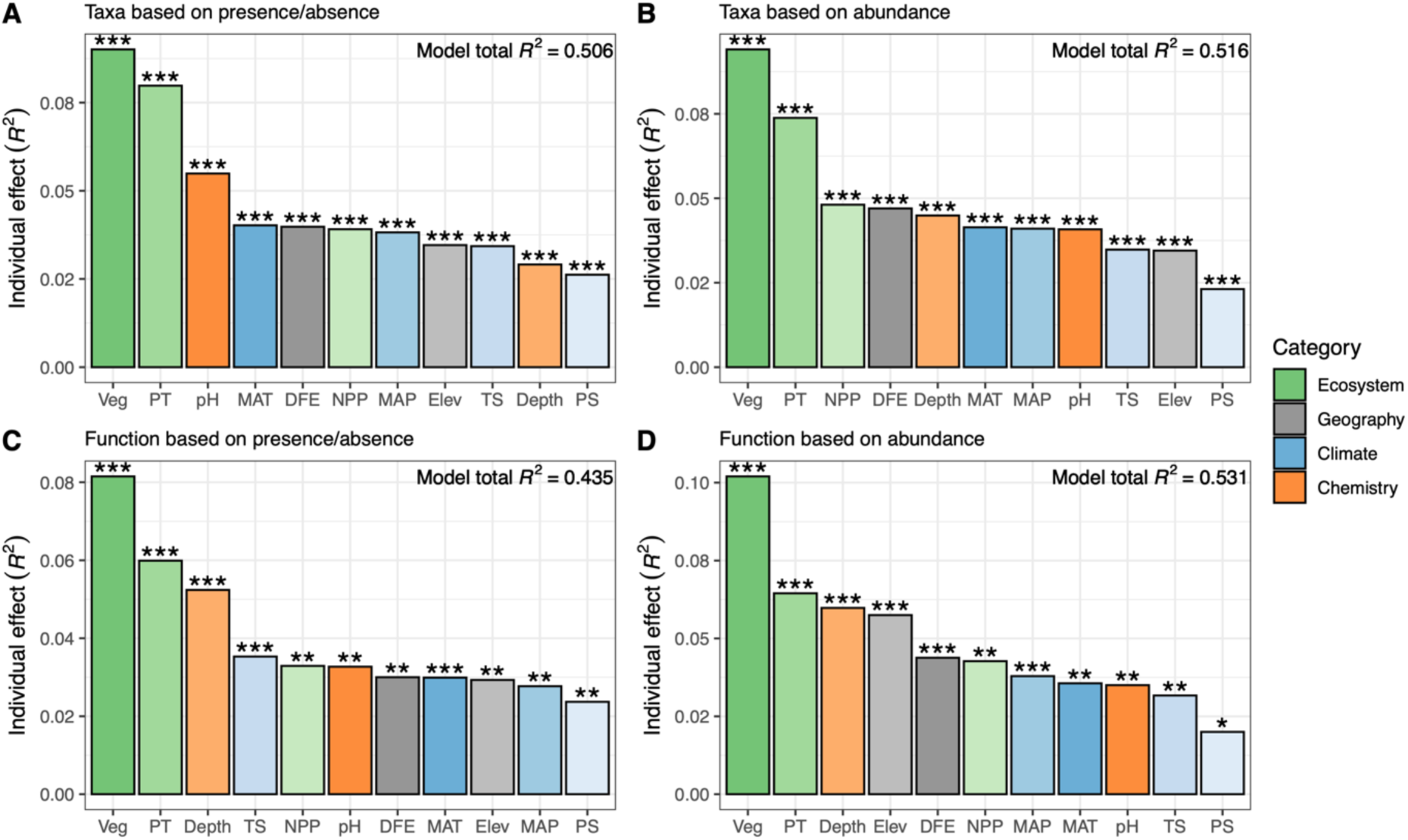
Ecological drivers of peatland microbiome taxonomy and function. Relative importance (adjusted *R*^2^ values) of individual predictor variables on the **(A, B)** taxonomic and **(C, D)** functional community membership and structure based on the **(A, C)** Jaccard (presence/absence) and **(B, D)** Bray–Curtis (abundance) distance metrics, respectively, by variation and hierarchical partitioning. The analysis was based on a subset with available pH data (n = 88). Ecological variables are coloured in gradients based on the category they belonged to. Also shown on each plot is the adjusted *R*^2^ value of the total model. Veg, dominant vegetation type; PT, peatland type; MAT, mean annual temperature; DFE, distance from the equator; NPP, net primary productivity; MAP, mean annual precipitation; Elev, elevation; TS, temperature seasonality; PS, precipitation seasonality. \**p* < 0.05, \*\**p* < 0.01, \*\*\**p* < 0.001.

### Depth and pH exert contrasting effects on peatland taxonomic and functional community membership

Depth was also found to be an important driver of peatland microbiome function, ranking third just after dominant vegetation and peatland type and explaining more than 5.2% of the variation; however, surprisingly, it ranked second to last in explaining taxonomic community membership, accounting for only 2.9% of the variation (Fig. 3). In contrast, pH played a prominent role in controlling taxonomic community membership, ranking third and explaining 5.5% of the variation, but it had the third lowest ranking and exhibited a smaller effect (3.5%) on the functional community structure. The predominant role of pH on the microbial community composition agrees with previous research on individual peatlands and the global topsoil microbiome[11, 29, 30], and may be attributed to the direct effect of pH or related variables, such as cation concentrations[31]. Depth has been identified as a significant factor influencing peatland microbial composition[11, 17]; however, these studies did not examine its relative effect compared to pH. Our finding of a larger effect of pH compared with depth on the taxonomic membership of the peatland microbiome suggests that the latter exerts a stronger control on some specific taxa, such as methanogens, but not the others. However, the effect of depth may have been underestimated here due to the underrepresentation of metagenomes collected at depths below 100 cm. Alternatively, the sampling resolution within the top 50 cm may have been insufficient, particularly around the water table, where shifts in oxygen availability can drive pronounced changes in microbial community composition and function.

Among the other predictor variables, elevation also appeared to be a strong driver of functional community structure of peatlands (ranking fourth and explaining 5.8% of the variation), although its effect on taxonomy was small (ranking fourth lowest or lower). Climate factors, in particular mean annual temperature (MAT) and mean annual precipitation (MAP), also played a role in controlling peatland community taxonomy and function. However, their effects were not major as they consistently ranked in the lower half of the rankings and explained at most 4.1% of the variation.

### Different peatland types host distinct microbiomes

We then characterised the taxonomic composition of the peatland microbiome. Here, to capture the vast taxonomic diversity of unknown and uncultivated microbial species provided by metagenome-assembled genomes (MAGs), we used MetaPhlAn 4[32] for taxonomic profiling of the metagenomes and mapped the resulting profiles to the Genome Taxonomy Database (GTDB) taxonomy (r220)[33]. Under the GTDB framework, alphanumeric nonstandard placeholder names are given to groups lacking cultured representatives based on a set of nomenclatural rules[34]. To preserve literature continuity, former names or synonyms were also included here when appropriate.

### Northern bog microbiome

At the phylum level, the northern bog microbiome was dominated by *Acidobacteriota* (28.3%), *Actinomycetota* (17.4%), *Pseudomonadota* (formerly *Proteobacteria*, 12.0%), and *Chloroflexota* (9.7%). At the family level, the uncultivated families UBA7540 (*Acidobacteriota*, 18.7%), RAAP-2 (*Actinomycetota*, 12.5%), UBA8260 (*Chloroflexota*, 8.5%), and Bog-38 (*Halobacteriota*, 7.0%) were the most abundant (Fig. 4, Supplementary Fig. 5). All of these families are commonly found in acidic environments. Differential abundance analysis (DAA) using both ANCOM-BC2 and MaAsLin 3 confirmed the higher abundance of UBA7540 and RAAP-2 in northern bogs compared with fens, UBA8260 in northern bogs compared with fens and TPs, and Bog-38 in northern bogs compared with TPs (Fig. 5B, Supplementary Table 3).

**Fig. 4.**
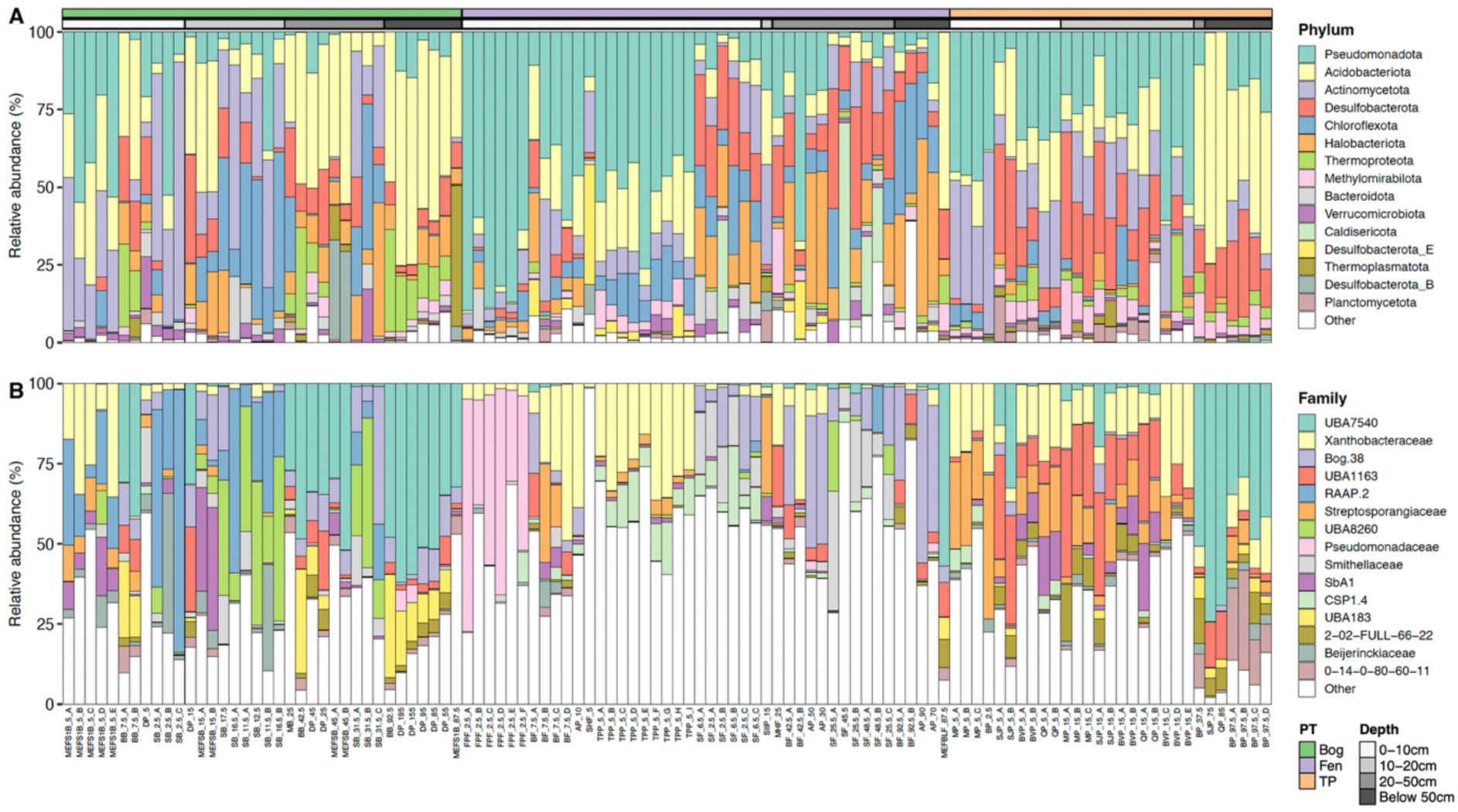
Taxonomic composition of the peatland microbiome. Stacked bar plots of the top 15 most abundant microbial **(A)** phyla and **(B)** families across the 109 metagenomes. Metagenomes are named using the syntax site_depth_replicate and annotated according to the peatland type (PT) and sampling depth. TP, tropical peatland.

**Fig. 5.**
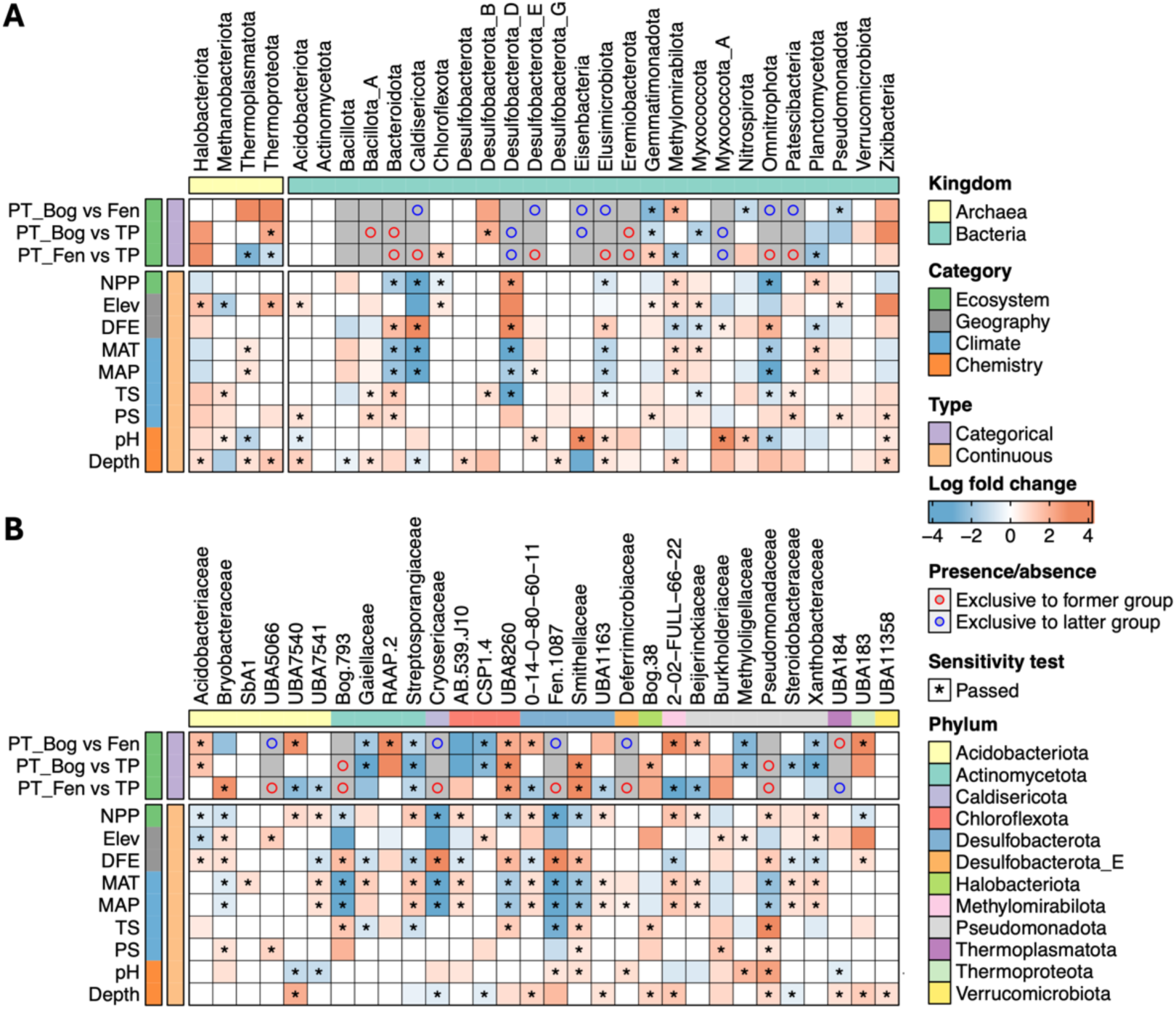
Impact of ecological variables on the abundance of peatland microbial taxa. Heatmaps showing the log fold changes in absolute abundance of the top 30 most abundant microbial **(A)** phyla and **(B)** families in relation to ecological variables as identified by ANCOM-BC2. Only taxa with a prevalence > 10% are shown. Each cell is coloured to represent significant changes in absolute abundance: red indicates increased abundance in the former as compared with the latter (reference) group for categorical variables and increased abundance per unit increase of continuous variables, whereas blue indicates the opposite. Entries marked with an asterisk have successfully passed the sensitivity analysis for pseudo-count addition. Cells in grey indicate the presence of structural zeros in at least one group of comparisons. Red and blue circles denote the exclusive presence in the former and latter group, respectively. Continuous variables were standardised before the calculations. Results of pH were based on a subset with available pH data (n = 88). Ecological variables are annotated according to the category and variable type they belonged to, whereas microbial taxa are annotated according to the kingdom or family they belonged to. Refer to Fig. 3 for the abbreviations of the variables. PT, peatland type; TP, tropical peatland.

*Acidobacteriota* is a ubiquitous group of large polysaccharide degraders in acidic peatlands and has been shown to contain a large genetic repertoire to degrade *Sphagnum* cell wall polysaccharides under both aerobic and anaerobic conditions[35]. The family UBA7540 within the phylum belongs to the former *Acidobacteria* subdivision 1[10], and consists of potentially facultative anaerobic sulphate reducing bacteria (SRB) that perform dissimilatory sulphate reduction or fermentation under anoxic conditions and can switch to aerobic respiration using plant-derived polysaccharides or low-molecular weight organic compounds under oxic conditions[36]. Given the generally low sulphate levels in bogs and the dominance of this family in deeper peats in our dataset (Fig. 5B, Supplementary Fig. 5B, Supplementary Table 3), these UBA7540 members presumably primarily perform fermentation rather than the other two processes in northern bogs. *Actinomycetota* and the *Chloroflexota* family UBA8260 (synonym *Ca. Aeolococcaceae*) are also known to aerobically degrade plant-derived OM[37, 38]. These collectively highlight the significant role of the northern bog microbiome in aerobic and anaerobic degradation of plant-derived OM.

*Halobacteriota*, previously part of *Euryarchaeota*, was the most abundant archaeal phylum recovered in this study. Within this phylum, the family Bog-38 (also known as *Ca.* Methanoflorentaceae or Rice Cluster II) consists of obligate hydrogenotrophic methanogens widespread in high methane-flux habitats, including rich paddies and anoxic freshwater sediments, and is thus believed to be a major contributor to global methane production[39]. In fact, Bog-38 members were not only abundant (7.0%) and highly prevalent in northern bogs (detected in 86% of metagenomes and 100% of sites) but also abundant (9.0%) and moderately prevalent in northern fens (detected in 59% of metagenomes and 67% of sites), albeit less abundant (0.8%) in TPs (Supplementary Fig. 5B, 6A). In comparison, canonical methanogenic classes detected here, including *Methanocellia*, *Methanomicrobia*, *Methanosarcinia*, and *Methanobacteria*, generally occurred in lower abundance, especially in bogs (Supplementary Fig. 6A). These results indicate that Bog-38 members are likely key methanogens in northern bogs and fens but not in tropical peatlands, despite higher methane fluxes in these systems. Their lesser importance in northern fens and especially TPs likely reflects alternative pathways of methanogenesis in those settings and is discussed below.

### Northern fen microbiome

At the phylum level, the northern fen microbiome was dominated by *Pseudomonadota* (31.3%), *Halobacteriota* (11.2%), *Acidobacteriota* (10.8%), and *Desulfobacterota* (10.4%). At the family level, *Xanthobacteraceae* (*Pseudomonadota*, 11.8%), Bog-38 (9.0%), *Pseudomonadaceae* (*Pseudomonadota*, 6.9%), *Smithellaceae* (*Desulfobacterota*, 4.9%), and CSP1-4 (*Chloroflexota*, 4.3%) were the most abundant. DAA confirmed the higher abundance *Xanthobacteraceae* and CSP1-4 in northern fens compared with bogs, as well as *Smithellaceae* compared with TPs (Fig. 5, Supplementary Table 3).

*Xanthobacteraceae* in northern fens was dominated by the genus *Pseudolabrys* (8.6%), which has been shown to encode a *Nitrobacter*-like *nxrA* and could represent a new nitrite oxidising bacteria (NOB) or incomplete denitrifier[40]. This, together with the fact that members of the uncultivated *Chloroflexota* family CSP1-4 can also perform denitrification[41], underscore a prominent role of the northern fen microbiome in denitrification/nitrification. *Desulfobacterota*, formerly part of *Deltaproteobacteria*, consists of obligate anaerobic SRB. However, instead of dissimilatory sulphate reduction, members from *Smithellaceae* are likely involved in syntrophic interactions, providing fermentation products to methanogens, such as Bog-38 which was abundant here[42].

### Tropical peatland microbiome

At the phylum level, the TP microbiome was dominated by *Pseudomonadota* (26.2%), *Acidobacteriota* (21.9%), *Desulfobacterota* (17.0%), and *Actinomycetota* (16.4%). At the family level, UBA7540 (15.0%), *Xanthobacteraceae* (14.5%), UBA1163 (*Desulfobacterota*, 12.6%), *Streptosporangiaceae* (*Actinomycetota*, 10.7%), and 2-02-FULL-66-22 (*Methylomirabilota*, 4.6%) were the most abundant. DAA confirmed the higher abundance of UBA7540, UBA1163 and 2-02-FULL-66-22 in TPs compared with northern fens, *Xanthobacteraceae* in TPs compared with northern bogs, and *Streptosporangiaceae* in TPs compared with both northern bogs and fens (Fig. 5B, Supplementary Table 3).

The uncultivated family UBA1163 (class BSN033) belongs to a group (cluster F) which has the second highest sulphur cycling potential within *Desulfobacterota*, just after *Desulfovibrionia[42]*. Together with potential SRB from UBA7540, the TP microbiome, at least at sites included in this study, appears to be dominated by microbes with sulphur reducing capabilities. Similar to northern fens, *Xanthobacteraceae* in TPs was dominated by the genera *Pseudolabrys* (6.6%) and BOG-931 (5.9%), both of which are potentially NOB or incomplete denitrifiers[40]. The uncultivated family 2-02-FULL-66-22 belongs to the order *Methylomirabilales* within *Methylomirabilota* (synonym candidate division NC10), which contains nitrite-dependent anaerobic methane oxidising (N-DAMO) bacteria that couple anaerobic methane oxidation to nitrite reduction[43]. They exhibited a higher abundance than canonical methanotrophs from the classes *Alphaproteobacteria* and *Gammaproteobacteria* here (Supplementary Fig. 6B), indicating their roles as important methanotrophs in at least these TPs. All together, these dominant taxa highlight the wide role of the TP microbiome in dissimilatory sulphate reduction, denitrification/nitrification, and anaerobic methanotrophy.

### The metabolic potential for plant OM degradation is higher in northern bogs and at greater depths

We then investigated the metabolic potential of the peatland microbiome for OM degradation by characterising carbohydrate-active enzymes (CAZymes) in the metagenomes. CAZyme genes involved in the degradation of major plant cell wall components, including cellulose, hemicellulose, pectin, and lignin, were detected in all peatland metagenomes (Fig. 6). However, DAA revealed a higher abundance of most of them (77.8%) in northern bogs compared with TPs, and some of them (33.3%) in northern bogs than in fens, with lignin being a notable exception, indicating a higher metabolic potential of the northern bog microbiome for plant OM degradation (Fig. 6, Supplementary Table 4). This finding coincides with, and expands upon, previous research reporting a higher OM degradation potential of the microbiome in the ombrotrophic nutrient-poor site compared with the minerotrophic nutrient-rich site of a tropical peatland[18], and is consistent with isotopic evidence that bacteria in bogs incorporate greater proportions of carbohydrate OM than those in other settings[44] or in tropical peatland[45]. The higher potential for plant OM degradation in northern bogs suggests that these peatland ecosystems could become more important sources of CO₂ and CH_4_ emission due to increased OM degradation under climate warming, especially given that bogs tend to accumulate more C than fens[46]. However, previous research suggests that viral predators and/or metabolic capacity overlap within the bog microbiome could prevent efficient OM degradation despite their high metabolic potential[11]. Moreover, microbial activity in bogs could be inhibited by organic compounds produced by *Sphagnum* mosses, the dominant vegetation in many bogs, including polyphenols such as sphagnum acid and the cell wall polysaccharide sphagnan, which contributes to environmental acidification[47].

**Fig. 6.**
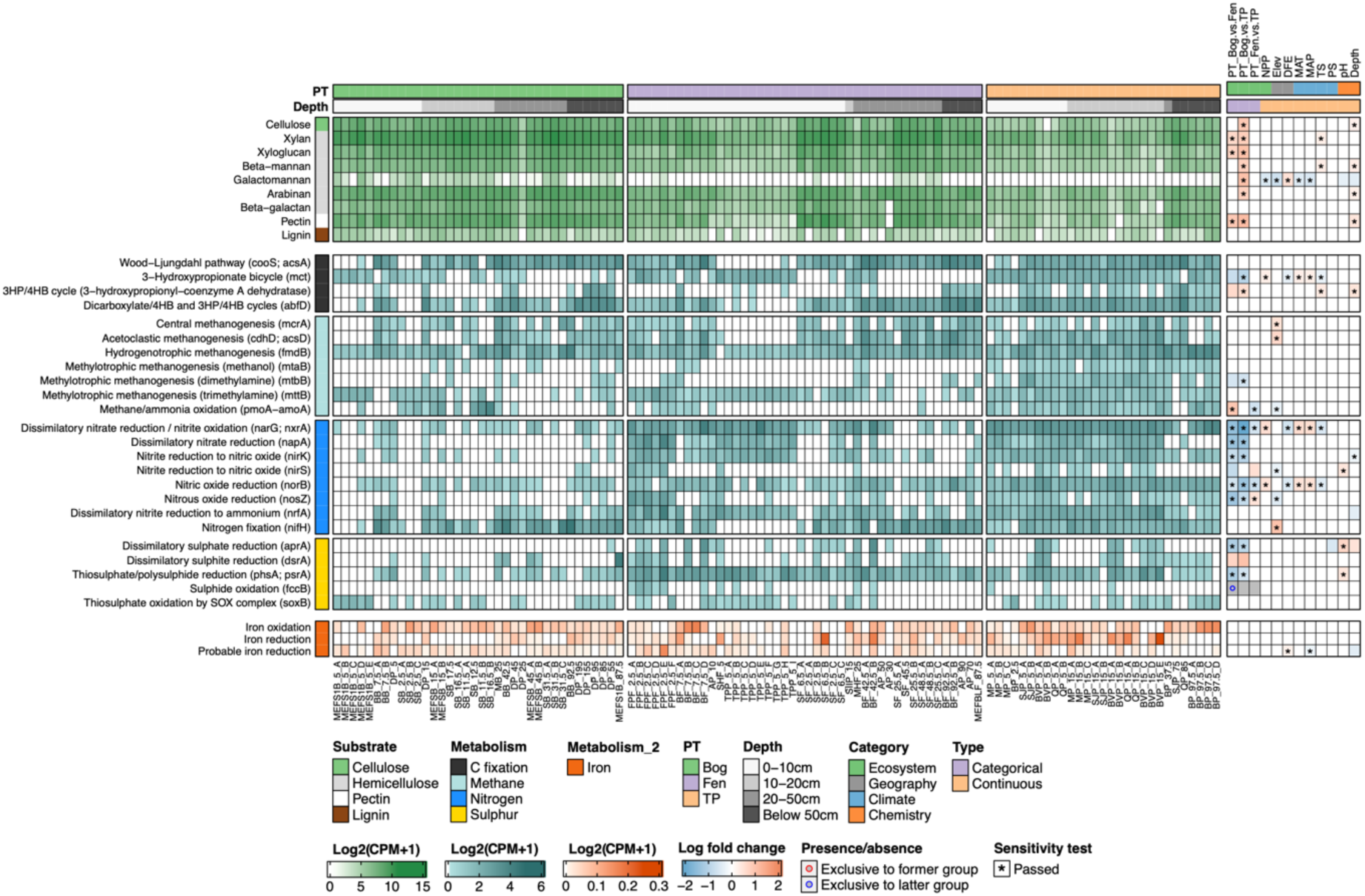
Metabolic potential of the peatland microbiome for key biogeochemical processes. Shown in the main heatmap are the relative abundances of plant cell wall-degrading CAZyme genes aggregated by CAZyme substrates (upper panel), marker genes for carbon fixation, methane, nitrogen, and sulphur cycling (middle panel), and iron cycling genes aggregated by function (lower panel) in log2-transformed copies per million (CPM) across the 109 metagenomes. A pseudo-count of 1 was added before log transformation to circumvent the zero-inflation problem. Heatmap to the right shows the log fold changes in absolute abundance of the genes in relation to different ecological variables as identified by ANCOM-BC2. Refer to Fig. 5 for annotations of the heatmap. Genes are annotated according to the substrates the CAZymes act on or metabolisms they involve in; metagenomes are annotated according to the peatland type (PT) and sampling depth; and ecological variables are annotated according to the category and variable type they belonged to. Refer to Fig. 3 for the abbreviations of the variables. TP, tropical peatland.

CAZyme genes for the degradation of cellulose, the hemicelluloses beta-mannan and arabinan, and pectin were found to be more abundant in deeper peats, observations also reported in other peatland and permafrost soil[17, 48]. This trend aligns with the higher abundance of the dominant phylum *Acidobacteriota* and family UBA7540 in northern bogs and TPs at depth (Fig. 4, 5, Supplementary Fig. 5, Supplementary Tables 3 and 5), reinforcing their key role in plant OM degradation in these peatland ecosystems, especially at depth. However, our dataset contained few metagenomes from greater than 100 cm depth, where reactive pools of OM are often depleted and alternative pathways of OM degradation, such as fermentation, could dominate[17].

Intriguingly, the metabolic capacity for lignin degradation did not differ among peatland types nor with depth, despite lignin being absent in *Sphagnum* moss and abundant in fen and TP vegetation[49] as well as in deeper peat layers[50]. The lack of significant depth-related differences in lignin-degrading capacity may be partly attributed to the smaller sample size at greater depths (>50 cm), which inherently reduces the statistical power required to detect true biological signals across the vertical gradient. This pattern may also reflect the presence of novel lignin-degrading enzymes, such as methyltransferase 1 (MT1, *mtgB*) in *Bathyarchaeia[51]*, that are not included in commonly used CAZyme databases.

### The tropical peatland microbiome exhibits a broad metabolic potential for key biogeochemical processes across all depths

We continued to investigate the metabolic potential for other key peatland biogeochemical processes by analysing selected marker genes (Fig. 6, Supplementary Table 6). Intriguingly, we observed a broad metabolic potential for most of these processes in TPs. These included the Wood-Ljungdahl (WL) (prevalence of *cooS; acsA*: 93.1%) and 3-hydroxypropionate (3HP) (*mct*: 96.6%) carbon fixation pathways, methanogenesis (*mcrA*: 65.5%), methane/ammonia oxidation (*pmoA-amoA*: 89.7%), denitrification (*narG; nxrA*: 96.6%, *napA*: 79.3%, *nirK*/*nirS*: 96.6%, *norB*: 100%, and *nosZ*: 86.2%), dissimilatory nitrate reduction to ammonium (DNRA) (*nrfA*: 96.6%), nitrogen fixation (*nifH*: 96.6%), thiosulphate/polysulphide reduction (*phsA; psrA*: 86.2%), iron oxidation (82.8%), and iron reduction (79.3%). Importantly, the potential for these processes was consistently present across depths, although the smaller sample size at greater depths should be noted. By contrast, for instance, methane/ammonia oxidation tended to be more prevalent in shallower peats in northern bogs and fens. The greater importance of the 3HP pathway, methane/ammonia oxidation, denitrification, and thiosulphate/polysulphide reduction in TPs compared with northern bogs and/or fens was further supported by DAA (Fig. 6, Supplementary Table 7). Notably, while the prevalence and abundance of *mcrA* was comparable across peatland types (prevalence in northern bog: 61.1%, northern fen: 61.4%, TP: 65.5%), 94.7% of the TP metagenomes harbouring the *mcrA* gene exhibited the potential for all three types of methanogenesis, compared to only 45.5% of northern bogs (Fisher’s exact test, *p* < 0.001) and 77.8% of northern fens (*p* > 0.05), suggesting that the TP microbiome exhibits a greater methanogenesis potential using a wider range of substrates, at least compared to northern bogs.

In contrast, northern bogs in general showed a low, and frequently the lowest, prevalence of marker genes for these processes. These included methane/ammonia oxidation (38.9%), denitrification (*narG; nxrA*: 61.1%, *napA*: 16.7%, *nirK*/*nirS*: 33.3%, *norB*: 66.7%, and *nosZ*: 13.9%), DNRA (50.0%), dissimilatory sulphate reduction (*aprA*: 2.8%), dissimilatory sulphite reduction (*dsrA*: 19.4%), thiosulphate/polysulphide reduction (52.8%), and iron reduction (47.2%). The lower importance of denitrification, thiosulphate/polysulphide reduction, and dissimilatory sulphate reduction in northern bogs compared with northern fens and TPs were supported by DAA. The prevalence of the northern fen microbiome’s potential for these processes was often intermediate between that of northern bogs and TPs. However, it was highest for thiosulphate/polysulphide reduction (100%) and lowest for methane/ammonia oxidation (20.5%) and iron oxidation (63.6%).

### A notable proportion of northern fen metagenomes exhibits an impaired potential for complete denitrification

The metabolic potential of the northern fen microbiome for denitrification was prevalent (*narG; nxrA*: 84.1%, *napA*: 69.4%, *nirK*/*nirS*: 72.7%, *norB*: 88.6%, and *nosZ*: 38.6%). The greater importance of denitrification in northern fens compared with bogs was supported by the higher abundance of all key marker genes for the process (*narG; nxrA, napA*, *nirK*, *norB*, and *nosZ*) in their metagenomes, as revealed by DAA. Notably, the *nosZ* gene, a marker for the final step in denitrification, was not detected in 42.9% of the northern fen metagenomes that harboured a complete set of genes for the reduction of nitrate, nitrite, and nitric oxide, suggesting a moderately high prevalence of the potential for incomplete denitrification in northern fens. In comparison, only 10.7% of the TP counterparts lacked *nosZ* (*p* < 0.05). This finding is corroborated by the higher prevalence of *nirK* than *nirS* in the northern fen metagenomes (72.7% vs 18.2%, *p* < 0.001); denitrifiers of the former type are more likely to perform incomplete denitrification[52]. It also extends previous research reporting a higher relative abundance of the *nor* gene compared with the *nosZ* gene in a minerotrophic seasonally frozen northern peatland[15], suggesting that these peatland ecosystems could be more susceptible to becoming N_2_O sources under environmental or climate change than other peatland types, especially given that heterotrophic denitrification has been shown to be the major pathway of N_2_O production in peatlands[15]. However, it is important to emphasize that the non-detection of genes in the metagenomes could be due to various reasons, including insufficient sequencing coverage, rather than their genuine absence in the microbiome. Nonetheless, the non-detection of these genes in the metagenomes suggests that they are in low abundance, even if present, making the processes they encode less prominent.

### Concluding remarks

By leveraging a curated dataset of 109 publicly available metagenomes from 20 near-natural peatland sites worldwide, we identified dominant vegetation and peatland type as the strongest predictors of peatland prokaryotic microbiome taxonomy and function and showed that different peatland types host distinct microbiomes. Importantly, we revealed a key role of the northern bog microbiome in plant organic matter degradation, an impaired metabolic potential of the northern fen microbiome for complete denitrification, and a broad potential of the tropical peatland microbiome for key peatland biogeochemical processes across all depths. These distinct large-scale patterns observed in different peatland types indicate that climate change may differentially affect the functioning of different peatland ecosystems.

There are a few limitations to this study that can inform future research. First, there are data gaps in the geographic coverage of currently available peatland metagenomic resources, and consequently in our dataset. Second, relatively few samples in our dataset were collected from depths below 50 cm, which may have led to an underestimation of depth-related effects. Third, the categorisation of peatland types lacked sufficient granularity, limiting the ability to make detailed comparisons, for instance, between herbaceous and woody tropical peatlands. Lastly, there was a lack of reusable physico-chemical data besides pH from the primary studies, preventing the examination of their effects on the microbiome. Nonetheless, this study has provided key insights into the large-scale patterns and ecological drivers of the peatland prokaryotic microbiome and led to several intriguing hypotheses that warrant further investigation. Future work should aim to acquire metagenomes from underexplored peatland types and deep peats, preferably from previously unexplored geographical regions, accompanied by the measurement and reporting of relevant physico-chemical metadata. In addition, integrating metagenomics with complementary approaches such as metatranscriptomics and stable isotope tracing will be essential for linking microbial community composition and functional potential to in situ activity in peatland ecosystems.

## Materials and Methods

### Metagenome dataset

Primary research articles related to peatland metagenomics were retrieved by searching PubMed using the following criteria: (metagenom*[Title/Abstract]) AND (peat*[Title/Abstract] OR mire[Title/Abstract] OR bog[Title/Abstract] OR fen[Title/Abstract]). Only studies including near-natural sites were selected. In this study, near-natural peatland sites were defined, following the descriptions in the primary studies and their cited references, as sites with minimal human disturbance (without evidence of drainage, cultivation, or other anthropogenic alteration) and without any record of major wildfire events. These included samples collected from control sites or prior to the start of the SPRUCE whole-ecosystem warming experiment[12, 53]. To establish a high-quality dataset and minimise methodological bias, only metagenomes sequenced paired-end with Illumina NovaSeq, NextSeq, or HiSeq were retained. Additionally, to create a balanced dataset comprising truly independent samples, one set of representative metagenomes was randomly selected per site if multiple metagenomes were available for the same sites at the same depths but collected at different timepoints. A total of 109 peatland metagenomes were then downloaded from the NCBI Sequence Read Archive (SRA) using SRA Toolkit v3.0.0, with accessions obtained from the primary studies[9–13, 15–19, 53–57].

### Metadata compilation

In this study, peatlands were broadly categorised into three major types: northern ombrotrophic bogs, northern minerotrophic fens, and tropical peatlands. Peatlands were also categorised based on the dominant vegetation types: *Sphagnum* moss, sedges, trees, and mixed. Depth, as a continuous variable, was derived from the mean of the sampling depth range and further grouped into four categories: 0–10 cm, 10–20 cm, 20–50 cm, and below 50 cm. pH data were retrieved from the primary studies or references therein. Net primary productivity (NPP) data (MOD17A3HGF v061) averaged over the period 2010–2023 at a resolution of 500 m and digital elevation data (ASTGTM v003) at 1-second resolution were downloaded from NASA using AρρEEARS. Climate data including mean annual temperature (MAT), mean annual precipitation (MAP), temperature seasonality (TS), and precipitation seasonality (PS) (where larger values of seasonality represent greater variability) averaged over the period 1970–2000 were downloaded from WorldClim 2.1[58] at 30-second spatial resolution. Values for individual peatland sites were then extracted using latitude and longitude coordinates with the *geodata* (0.6-2) R package. Distance from the equator (DFE) was calculated using the *geosphere* (1.5-20) R package with the Vincenty (ellipsoid) method. Ecological variables were grouped into four categories: ecosystem (peatland type, dominant vegetation type, and NPP), geography (elevation and DFE), climate (MAT, MAP, TS, and PS), and chemistry (pH and depth). Depth was categorised as a chemical variable here because it often relates to the physicochemical properties (e.g. redox potential and oxygen availability) of peat[3].

### Taxonomic and functional profiling of metagenomes

Metagenomes were first quality-filtered using Trimmomatic v0.39[59] with the following parameters: ILLUMINACLIP:TruSeq3-PE-2.fa:2:30:10 LEADING:3 TRAILING:3 SLIDINGWINDOW:4:15 MINLEN:36. Quality-filtered metagenomes were then subsampled to the smallest sample size (12 million reads) using Seqtk v1.4 to minimise the impact of variation in sequencing effort, differing by as much as 31-fold in read number, on subsequent sample comparisons[60]. Taxonomic profiling was performed using MetaPhlAn 4.0.6[32] against the CHOCOPhlAnSGB database (vJun23) with the parameters --stat_q 0.1 --min_mapq_val -1 as recommended for environmental samples. The resulting species-level genome bin (SGB)-based taxonomic profiles were then converted to the GTDB taxonomy (r220)[33] using the *sgb_to_gtdb_profile.py* script included in the package.

Read-based functional profiling was performed using HUMAnN 3.8[61] against the CHOCOPhlAnSGB (vOct22) and UniRef50 databases (v201901b)[62] as recommended for environmental samples. Gene families normalised for gene length were regrouped into KEGG orthologs (KOs) using the *humann_regroup_table* script included in the package. Feature tables were imported into QIIME2 v2024.5[63] and destratified using the *q2-sapienns* plugin. The resulting tables were then imported into R using the *qiime2R* (0.99.6) R package for downstream statistical analyses.

For the annotation of carbohydrate active enzymes (CAZymes), their substrates, and iron-related genes, quality-filtered metagenomic reads were first assembled into contigs using metaSPAdes v3.15.5[64]. Genes were then predicted from contigs longer than 500 bp using Prodigal v2.6.3[65]. CAZymes were annotated using run_dbcan v4.0[66] based on three approaches: HMMER (v3.4) against dbCAN HMMdb (v12), HMMER against dbCAN-sub HMMdb (v12), and DIAMOND against the CAZy database (release 07262023)[67], and those predicted by ≥ 2 approaches were kept. Substrates of CAZymes were predicted by HMMER against dbCAN-sub HMMdb. Normalised abundances of CAZyme families and substrates were calculated following an associated protocol[68]. Iron-related genes were annotated and grouped into gene categories according to their functions using FeGenie v1.2[69].

### Statistical analyses

All statistical analyses and plotting were performed in R v4.4.0. All pH-related analyses were conducted based on a subset of 88 metagenomes with associated pH data. Statistical significance was defined at *p* < 0.05.

#### Alpha and beta diversity analysis

Feature tables were first rarefied to the smallest sample size using the *rrarefy* function in *vegan* (2.6-6.1)[70]. Unmapped and ungrouped gene families were removed before the calculations. For alpha diversity analysis, the Shannon diversity index and number of SGBs and KOs were calculated using the *diversity* function in *vegan*. Difference between groups of categorical variables was assessed by Kruskal–Wallis test followed by two-sided Dunn’s test using *rstatix* (0.7.2). The Holm–Bonferroni method was used for the correction of multiple comparisons. Correlations between alpha diversity indices and continuous variables were assessed by Spearman’s correlation and displayed in scatter plots using *ggpubr* (0.6.0). For beta diversity analysis, principal coordinates analysis (PCoA) based on the Jaccard and Bray–Curtis distance metrics was conducted using the *capscale* function in *vegan*. Permutational multivariate analysis of variance (PERMANOVA) was then performed using the *adonis2* function in the same package.

#### Drivers of microbiome taxonomy and function

The relative importance of individual predictor variables on the taxonomic and functional community membership and structure was assessed based on the Jaccard and Bray–Curtis distance metrics, respectively, using variation and hierarchical partitioning for canonical analysis with *rdacca.hp* (1.1-0)[25]. In this approach, variation partitioning was first used to estimate the unique and shared variation among predictor variables, followed by hierarchical partitioning to calculate the relative importance of individual variables across all possible model subsets without conditioning effects. This approach is particularly useful when correlation (multicollinearity) among variables is intermediate or strong[25].

#### Differential abundance analysis

Differential abundance analysis (DAA) for both categorical and continuous variables was conducted using the ANCOM-BC2 approach[71] with the *ancombc2* function in *ANCOM* (2.8.1). For MetaPhlAn taxonomic profiles, relative abundances were converted to count data by multiplying each value by the subsampled sequencing depth (i.e. 12 million reads), and the resulting counts were used as input to ANCOM-BC2. For UniRef50 gene families, CAZyme genes, and iron-related genes, read counts normalised for gene length in reads per kilobase (RPK) were used as input. Normalisation by gene sequence length is essential for accurate estimation of gene abundance[72, 73] and is not expected to have any major impact on the overall results[74]. ANCOM-BC2 estimates and corrects both sample-specific and feature-specific biases and is specifically designed for multigroup analyses by effectively controlling mixed directional false discovery rate. Continuous variables were first standardised before the analysis. Features having structural zeros, defined as being completely (or nearly completely) missing in one or more groups in comparisons, or present in less than 10% of all samples were removed from further analysis by ANCOM-BC2. For features having structural zeros, we defined an exclusive presence of them as being present in at least 20% of the samples in the non-structural zero group(s). The Holm–Bonferroni method was used for the correction of multiple comparisons. Changes in the prevalence of features were tested using two-sided Fisher’s exact tests using the *stats* package in base R. Significant associations with peatland type and depth identified by ANCOM-BC2 and discussed in the text were further validated using MaAsLin 3[75]. Relative abundances (for taxonomic profiles) and copies per million (CPM) values (for functional profiles) were used as input, and internal normalisation was therefore disabled.

## Data availability

All data analysed during the current study are publicly available. Accession numbers of all metagenomes included in this study are provided in Supplementary Table 1.

## Acknowledgements

This work was supported by a UKRI Advanced Research Grant (EP/X023214/1) to R.D.P. M.V. was supported by the NERC GW4+ Doctoral Training Partnership (NE/S007504/1). A.V.G-S. acknowledges support from the European Research Council (grant agreement no. 865403). We are grateful to all the authors who made their primary metagenome data publicly available and additionally thank Hongyan Wang for providing additional information about the Dongtu peatland. Computational work was carried out using the High Performance Computing facility of the Advanced Computing Research Centre at the University of Bristol.

## Author contributions

S.M.K.C. and C.B. conceived the study; S.M.K.C. compiled the datasets, performed the analyses, and wrote the original draft of the manuscript; R.D.P., M.V., A.V.G-S. and C.B. critically reviewed the manuscript. All authors approved the final version of the manuscript.

## Declarations

## Consent to Publish declaration

not applicable.

## Consent to Participate declaration

not applicable.

## Ethics declaration

not applicable.

**Correspondence and requests for materials** should be addressed to Simon Man Kit Cheung or Casey Bryce.

