## Supplementary figures 1 to 6 for "Structure and function of the peatland prokaryotic microbiome"

Simon Man Kit Cheung[1,*], Richard D. Pancost[1,2], Mike Vreeken[1,2], Angela V. Gallego-Sala[3], Casey Bryce[1,*]

[1]*School of Earth Sciences, Cabot Institute for the Environment, University of Bristol, Bristol, UK*

[2]*Organic Geochemistry Unit, School of Chemistry, University of Bristol, Bristol, UK*

[3]*Department of Geography, Faculty of Environment, Science and Economy, University of Exeter, Exeter, UK*

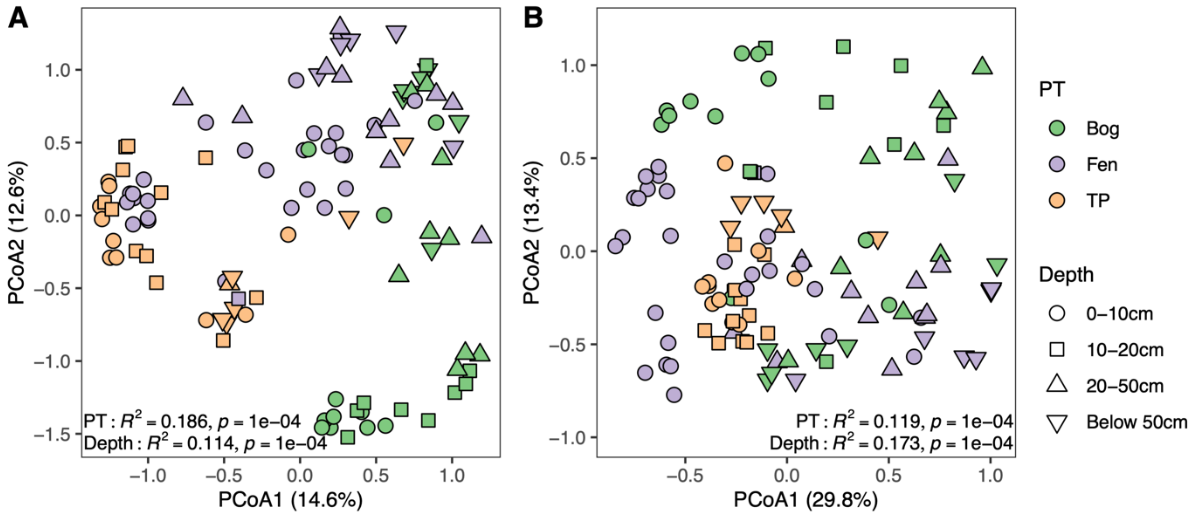


**Supplementary Fig. 1. Principal coordinates analysis (PCoA) plots of (A) taxonomic and (B) functional beta diversity based on the Jaccard (presence/absence) distance metric.** Values on the axes represent the percent variations explained. PERMANOVA statistics are provided on the plots. PT, peatland type; TP, tropical peatland.

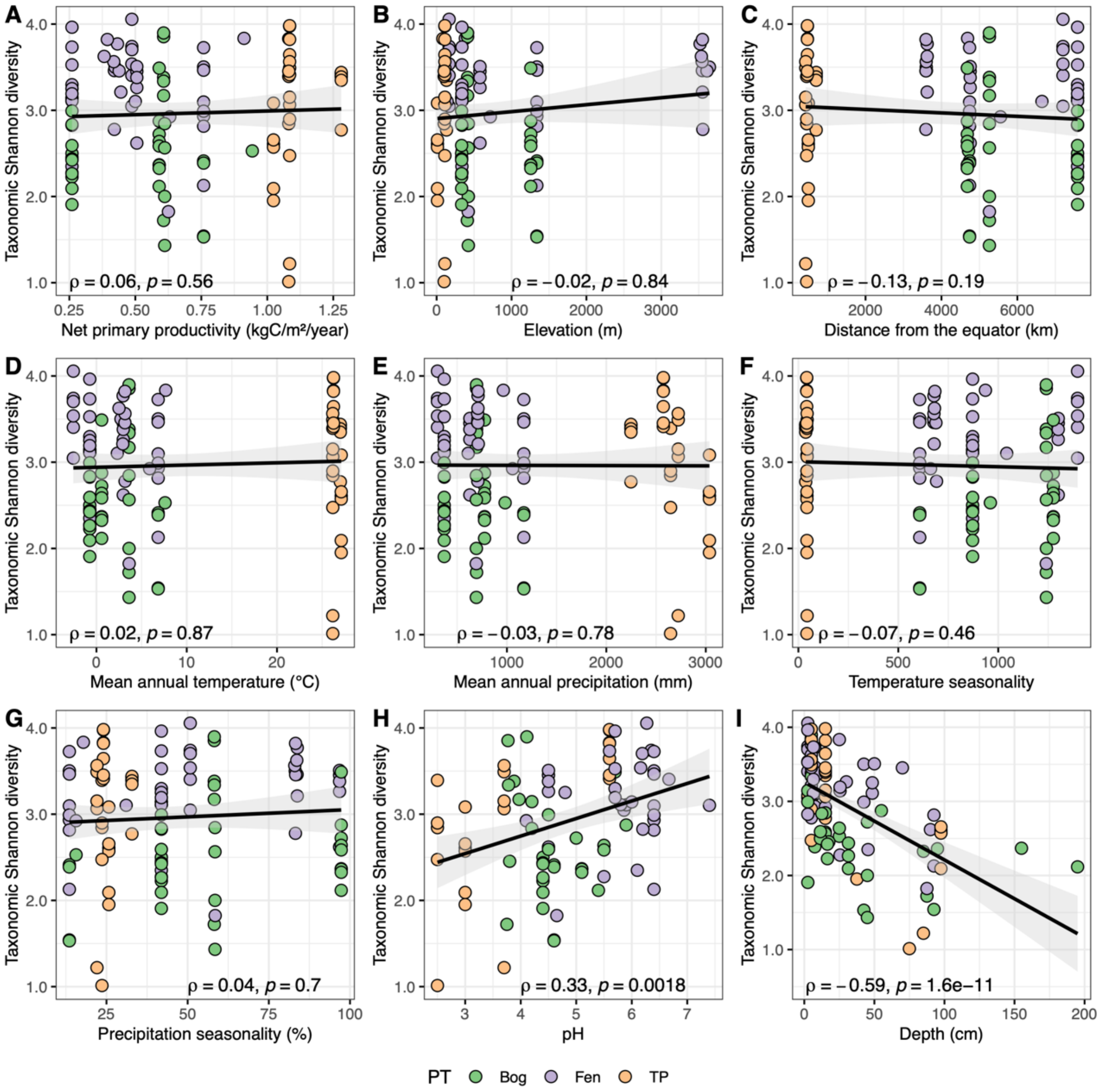


**Supplementary Fig. 2. Spearman's correlations between taxonomic alpha diversity and continuous variables.** Grey areas around the regression lines represent regions of 95% confidence interval. Panel H was based on a subset with available pH data (n = 88). PT, peatland type; TP, tropical peatland.

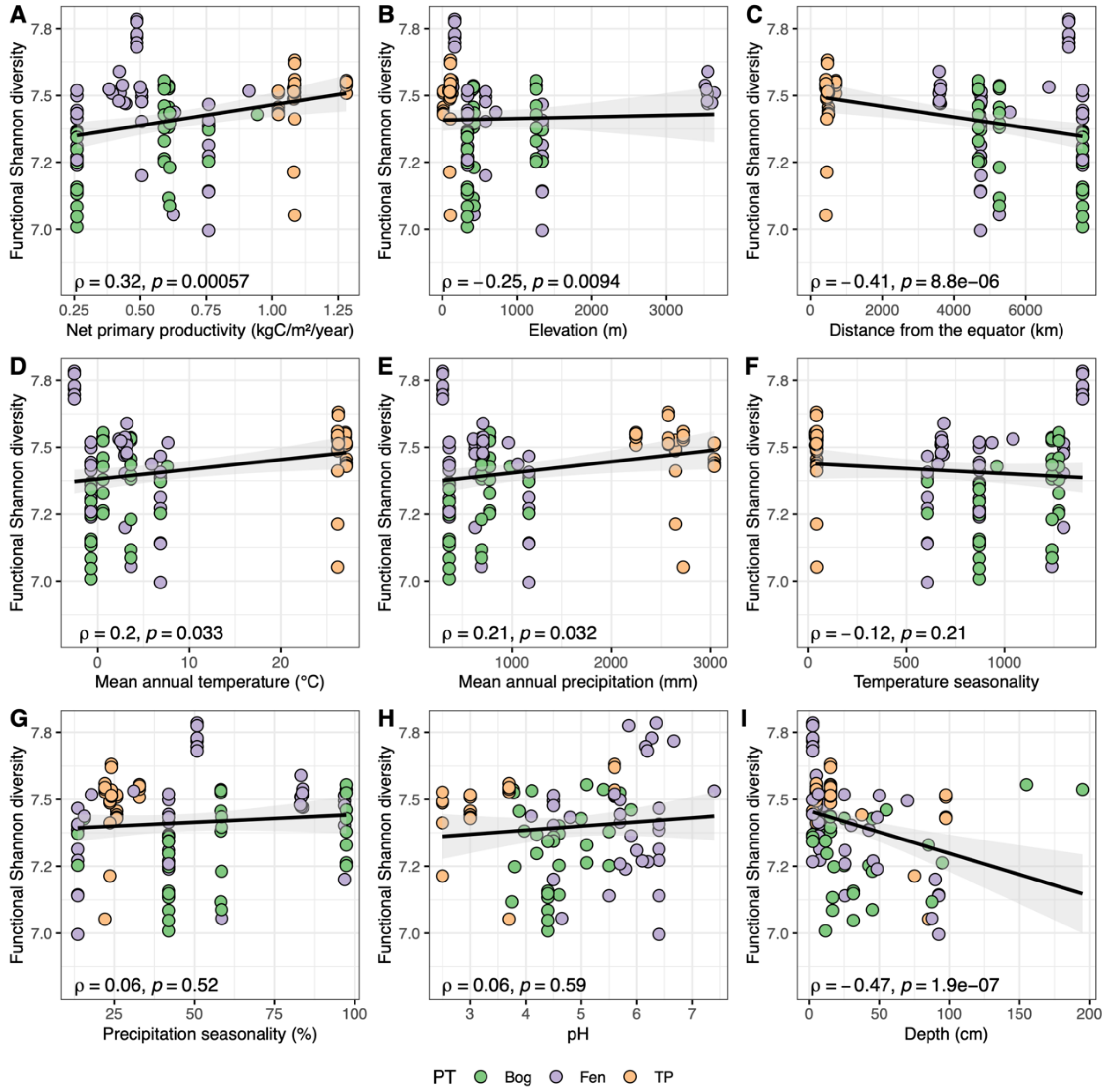


**Supplementary Fig. 3. Spearman's correlations between functional alpha diversity and continuous variables.** Grey areas around the regression lines represent regions of 95% confidence interval. Panel H was based on a subset with available pH data (n = 88). PT, peatland type; TP, tropical peatland.

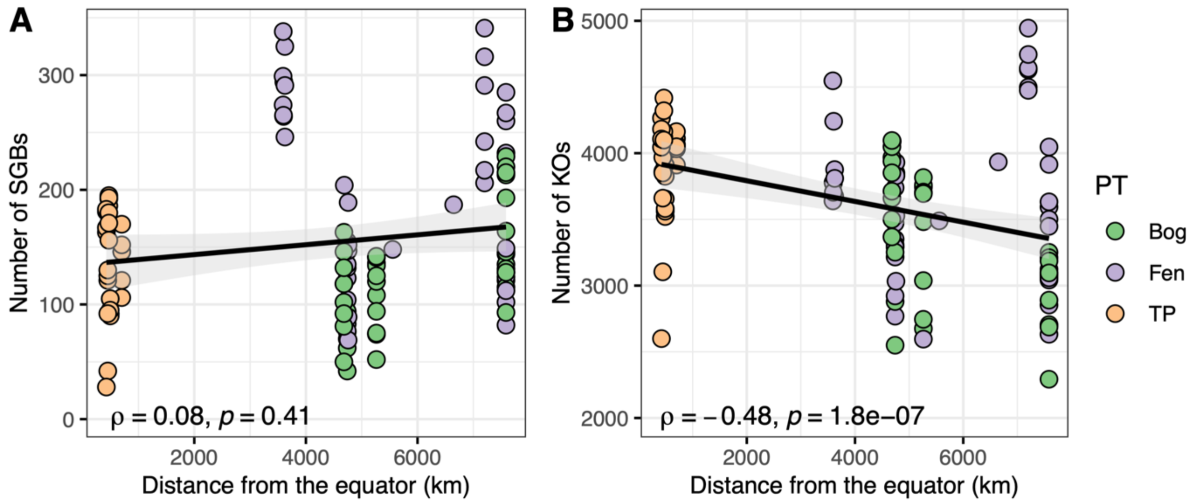


**Supplementary Fig. 4. Latitudinal diversity gradient analysis in taxa and function.** Spearman's correlations between distance from the equator and **(A)** the number of species-level genome bins (SGBs) and **(B)** the number of KEGG orthologs (KOs). Grey areas around the regression lines represent regions of 95% confidence interval. PT, peatland type; TP, tropical peatland.

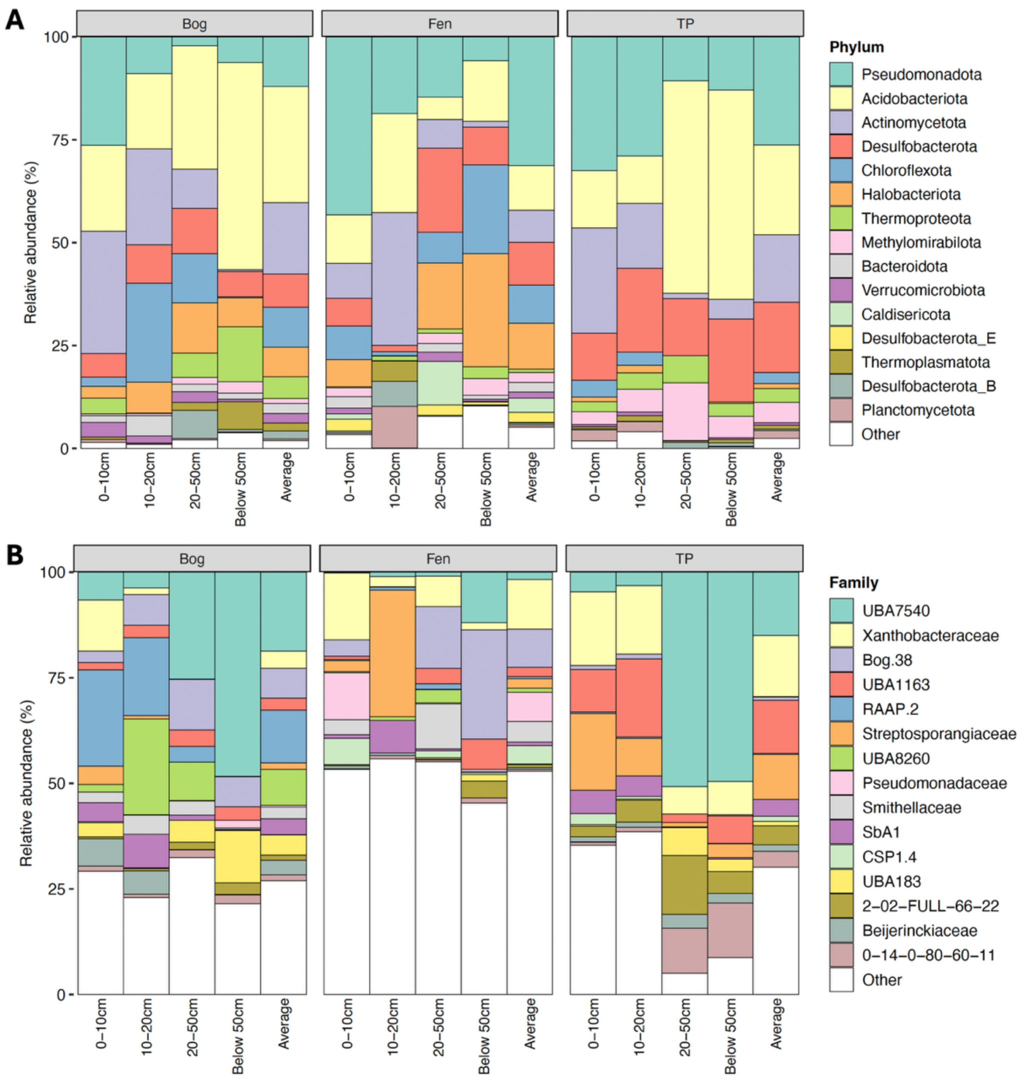


**Supplementary Fig. 5. Taxonomic composition of the peatland microbiome aggregated by peatland type and depth.** Stacked bar plots of the top 15 most abundant microbial **(A)** phyla and **(B)** families across depth in each peatland type. Also included in each facet is a bar representing the averaged values across all metagenomes from a particular peatland type. TP, tropical peatland.

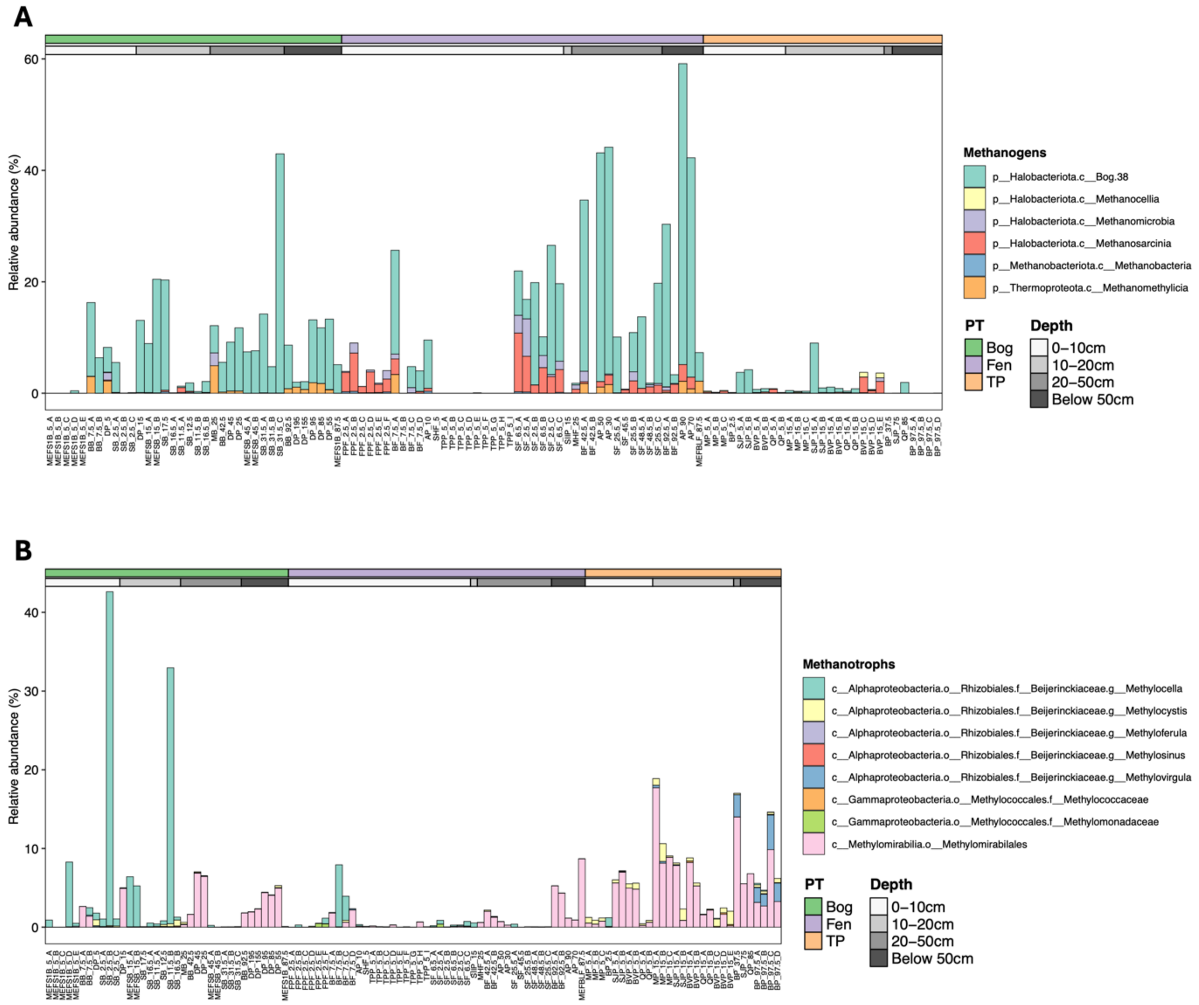


**Supplementary Fig. 6. Taxonomic composition of methanogens and methanotrophs.** Stacked bar plots of **(A)** methanogenic classes and **(B)** methanotrophs across the 109 metagenomes. Metagenomes are annotated according to the peatland type (PT) and sampling depth. TP, tropical peatland.
